# A Simple Enhancement for Gibson Isothermal Assembly

**DOI:** 10.1101/2020.06.14.150979

**Authors:** Brian A Rabe, Constance Cepko

## Abstract

Gibson Isothermal Assembly has become a widespread cloning method, with a multitude of advantages over traditional cut-and-paste cloning. It allows for scarless assembly of multiple fragments simultaneously and has become widely used for molecular cloning. We have found that a simple change to the formulation of the reaction mix, the addition of a single-stranded DNA binding protein, can substantially improve both the accuracy and efficiency of assembly, especially as the number of fragments being assembled increases. In addition, when creating this Enhanced Gibson Isothermal Assembly reaction mix in-house with homemade DNA ligase, the cost of the reaction can be reduced to less than $10 per milliliter.

## Introduction

For decades, assembling DNA constructs and libraries has been a mainstay of molecular biology. For much of that time, the traditional cut and paste methods were widely used, relying on restriction endonucleases and a DNA ligase to, for example, insert DNA sequences into a vector.^1^ The isothermal assembly method created by Gibson et al., often referred to as Gibson Assembly, marked an impressive advancement over cut and paste methods.^2^ Gibson Assembly allows for the simultaneous assembly of multiple DNA fragments based on overlapping terminal regions. It does not necessarily rely on restriction endonuclease recognition sites and is capable of scar-free assembly. It comes as no surprise, therefore, that this method has quickly become a popular method of choice for molecular cloning.

As described in Gibson et al., Gibson Assembly is an isothermal assembly reaction consisting of DNA fragments with homologous terminal regions and three enzymes and is run at an elevated temperature.^2^ T5 exonuclease is used to chew back the 5’ ends of the DNA fragments, leaving 3’ overhangs (Figure 1). The homologous regions on the 3’ ends can anneal, allowing for extension by a thermophilic DNA polymerase to fill the remaining gaps. Finally, a thermostable DNA ligase ligates the resulting nicks.

**Figure 1.**
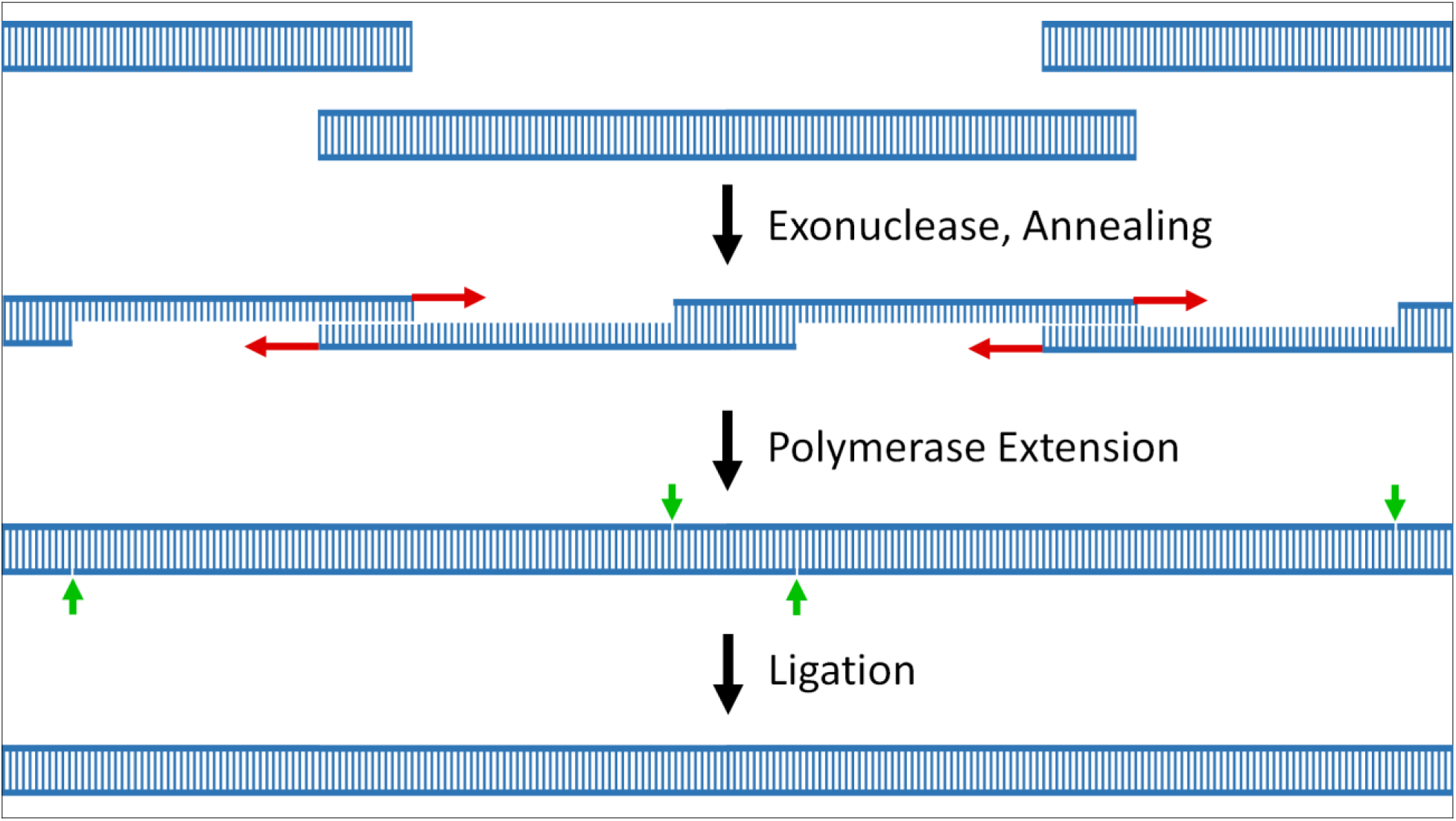
Schematic of Gibson Assembly. A brief schematic indicating the steps for isothermal assembly of DNA fragments as described by Gibson et al.^2^

While Gibson Assembly has proven invaluable, there is always a desire for even greater accuracy and efficiency. Interestingly, the T5 exonuclease also has a single-stranded endonuclease activity that may be detrimental to the reaction.^3^ This activity could cleave the single-stranded 3’ overhangs revealed by the exonuclease activity, resulting in undesired 3’ single-stranded DNA (ssDNA) ends (Figure 2a) which could assemble improperly or not at all. In order to address this problem, we added a fourth protein to the Gibson Assembly reaction, the single-stranded DNA-binding protein (SSB) from a thermophilic organism (ET SSB from New England Biolabs). SSB is known to bind ssDNA, protecting it from cleavage by endonucleases.^4^ Furthermore, SSB has been found to reduce secondary structure of ssDNA, improve PCR specificity, and improve the processivity of DNA polymerases.^5-8^ Thus, we hypothesized that ET SSB would protect the ssDNA ends from cleavage and might additionally improve the efficiency and specificity of these ends annealing, resulting in a more accurate and efficient Gibson Assembly reaction (Enhanced Formulation, Figure 2b).

**Figure 2.**
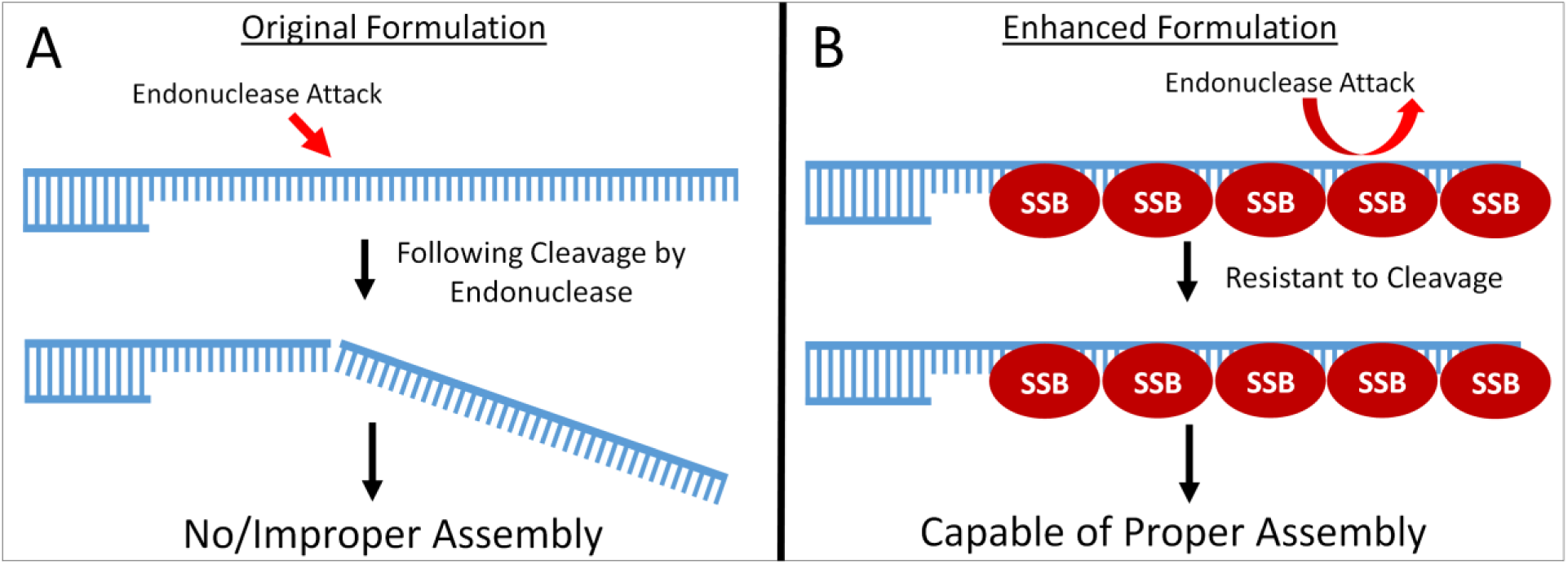
Schematic of Potential Mechanism of SSB. A – A depiction of the activity of T5 exonuclease’s ssDNA endonuclease activity on ssDNA ends created in a Gibson Assembly Reaction. B – A depiction of SSB’s ability to protect ssDNA ends created in a Gibson Assembly Reaction from endonuclease attack.

Strikingly, this formulation, which we refer to as Enhanced Gibson Assembly, is significantly more accurate and efficient than the original formulation and is comparable to or better than a commercially available high-fidelity assembly mix. Finally, by making this assembly mix in-house with homemade *Tth* DNA Ligase, the cost can be reduced dramatically.

## Results/Discussion

In order to test the accuracy and efficiency of various assembly mixes, we assembled fragments into the pUC19 vector that, when assembled, would result in bacterial expression of fluorescent proteins (Figure 3). We tested the assembly of two and six inserts. For the 2-insert assembly, the correctly assembled construct would result in Enhanced Green Fluorescent Protein (EGFP) expression (Figure 3a). For the 6-insert assembly, the correctly assembled construct would result in both EGFP and mCherry expression from a bicistronic transcript (Figure 3b). Thus, following transformation of these assembly reactions, the accuracy could be determined by scoring the percentage of resultant colonies that exhibited the correct fluorescence. In order to estimate the efficiency of the reaction, an intact plasmid containing a kanamycin resistance gene was included in each reaction as an internal transformation control.

**Figure 3.**
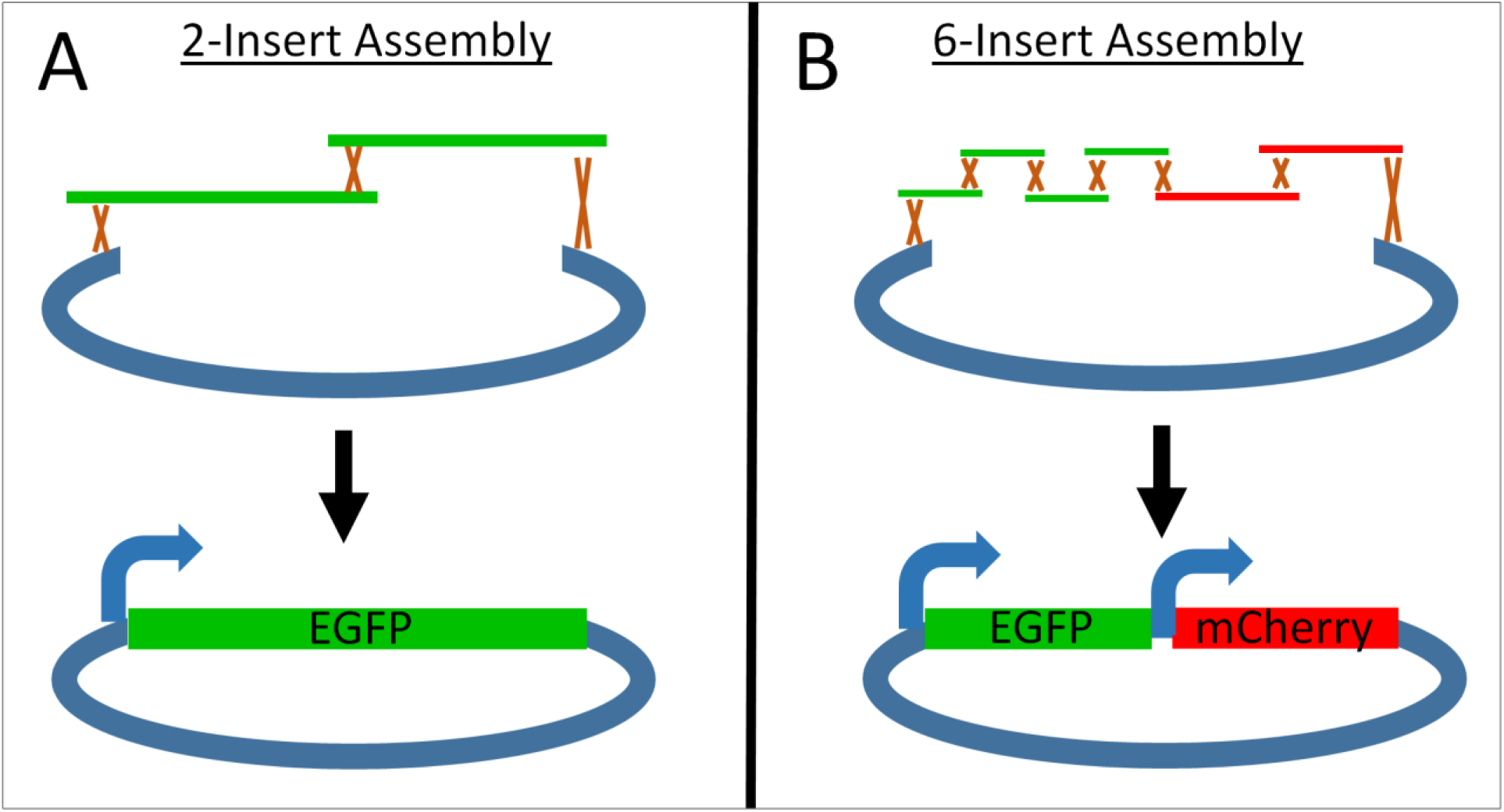
Schematic of Experimental Assembly Reactions. A – A 2-insert assembly creating a construct capable of expressing EGFP in transformed *E. coli*. B – A 6-insert assembling creating a construct capable of expressing both EGFP and mCherry from a bicystronic transcript in transformed *E. coli*.

**Figure 4.**
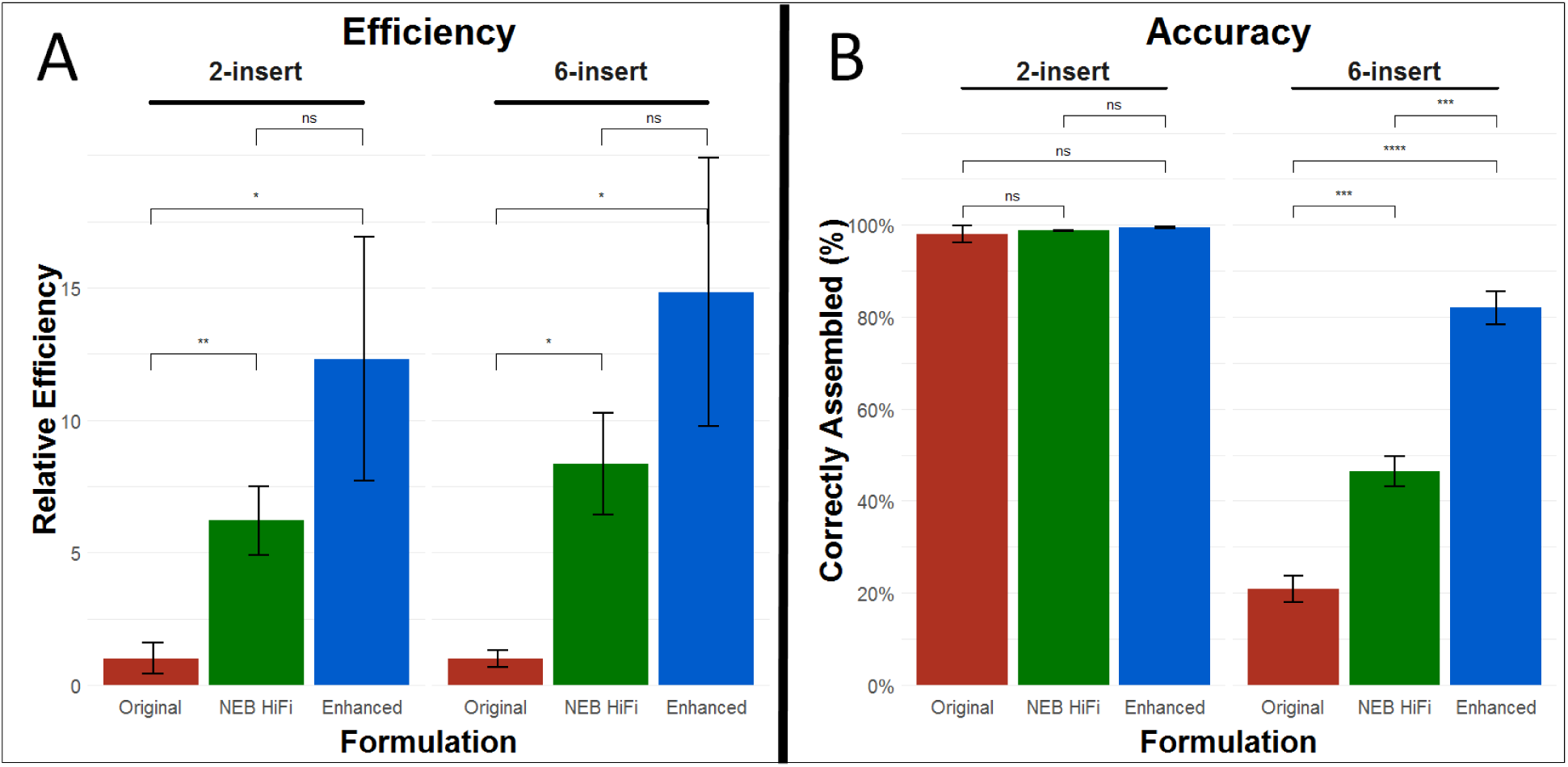
Quantification of Results of Assembly Reactions. A – The relative efficiency of each mix, normalized to the efficiency of the original formulation for both assembly schemes, n = 3 for each. B – The percent of transformants following assembly with each mix exhibiting correct fluorescence for both assembly schemes, n = 3 for each. A non-paired t-test was used for statistical analysis. Error bars represent standard error. ns - p>0.05; * - p <=0.05; ** - p<=0.01, *** - p<=0.001; **** - p<= 0.0001;

Compared to the original Gibson Assembly formulation, this enhanced formulation exhibited markedly increased accuracy for the 6-insert assembly (>80% compared to ∼20%, p = 3e-5), and greatly increased efficiency in both 2-insert and 6-insert assemblies (>12-fold increase, p < 0.05). Compared to the NEBuilder® HiFi DNA Assembly Master Mix (NEB E2621), the enhanced formulation showed a trend toward higher assembly efficiency, but this trend failed to reach statistical significance. However, the enhanced formulation showed significantly increased accuracy in the 6-insert assembly (>80% compared to ∼45%, p = 0.00024).

These results clearly demonstrate that the addition of ET SSB can increase the efficiency and accuracy of Gibson Assembly, especially as the number of fragments being assembled increases. While the exact mechanism has yet to be conclusively elucidated, it is possible that this ssDNA binding protein may facilitate specific hybridization and protect ssDNA overhangs from endonuclease attack.^4, 6^ The simplicity of this modification makes it easy to use in place of the original formulation without any other modifications. Furthermore, by making this mix in-house with homemade *Thermus thermophilus* DNA Ligase, this mix can be made for less than $10 per milliliter, making it accessible in a wide variety of research settings.

## Materials and Methods

Homemade Tth DNA Ligase – see supplementary protocol 1.

### Homemade Assembly Mixes

Both the original and enhanced Gibson Assembly mixes were based on recipes from *OpenWetWare*.^9^ Briefly, a 5x Isothermal Reaction Mix was created by combining 3 ml 1 M Tris-HCl, pH = 7.5 (ThermoFisher Scientific 15567027), 300 µl 1 M MgCl_2_ (Millipore Sigma M1028), 600 µl 10 mM dNTP mix (Goldbio D-900-1), 46.3 mg DTT (Goldbio DTT10), 20 mg NAD (Millipore Sigma N0632), 1.5 g PEG 8000 (Millipore Sigma 89510), and brought to 6 ml with UltraPure water (ThermoFisher Scientific 10977015). 320 µl aliquots were stored at -80C. To create a 1.33x Gibson Assembly mix, 40 units Phusion High-Fidelity DNA Polymerase (NEB M0530), 12 units T5 exonuclease (NEB M0363), and 6,400 units Tth DNA Ligase (homemade, supplementary protocol 1) were added to 320 ul of 5x Isothermal Reaction Mix. For the enhanced formulation, 10 ul ET SSB (500 µg/ml, NEB M2401S) was also added. UltraPure water (ThermoFisher Scientific 10977015) was added to bring the final volume of each to 1.2 ml. Gibson Assembly reaction mixes were dispensed into single use aliquots and stored at -20C.

### Preparation of DNA Fragments

All fragments were generated with PCR using Q5 Hot Start High-Fidelity DNA Polymerase (NEB M0493) and gel purified using the QIAquick Gel Extraction Kit (Qiagen 28704) using manufacturer protocols. Fragments containing portions of the EGFP or mCherry ORFs were cloned from plasmids containing these sequences, and the linear pUC19 vector was amplified from a circular pUC19 template (see supplementary table 1).

### Assembly Reactions

All reactions included each insert fragment at a 6-fold molar excess to the vector fragment. Each 10 µl assembly reaction included 4 fmol of vector along with 24 fmol of each insert fragment and 0.4 fmol of an intact supercoiled kanamycin-resistant plasmid (as a transformation control). For both the original and enhanced Gibson Assembly formulations, 2.5 µl of this DNA mixture was added to 7.5 µl of 1.33x master mix on ice. For the NEBuilder® HiFi reactions, 2.5 µl of this DNA mixture and 2.5 µl of UltraPure water was added to 5 µl of NEBuilder® HiFi DNA Assembly Master Mix (NEB E2621) on ice. After mixing, the reactions were moved directly to 50C for one hour and then placed back on ice. Both the 2-insert and 6-insert assemblies were run in triplicate for each reaction formulation.

### Transformation

15 µl of UltraPure water was added to each assembly reaction on ice and 2.5 µl of these diluted reactions were then transformed into 50 µl chemically competent DH5α cells (prepared in house, Zymo T3001) on ice. Following a five minute incubation on ice, 450 µl of pre-warmed SOC outgrowth medium (NEB B9020S) was added and the transformation was incubated at 37C with shaking (250 RPM) for one hour. 200 µl of each transformation was then plated on an LB-agar plate with carbenicillin (100 µg/ml) and another 200 µl was plated on an LB-agar plate with kanamycin (50 µg/ml). Plates were incubated overnight at 37C.

### Colony Counts and Analysis

Following overnight growth, total colonies on all plates were counted. For each LB-agar plate with carbenicillin, the number of colonies exhibiting correct fluorescence were counted (EGFP+ for 2-insert assembly, EGFP+/mCherry+ for 6-insert assembly) using a fluorescence stereoscope (Leica M165 FC). Accuracy was calculated as the percentage of colonies exhibiting correct fluorescence. Relative efficiency was a measure of the ratio between total carbenicillin resistant colonies to kanamycin resistant colonies for each transformation, normalized to the mean ratios for the original Gibson Assembly formulation assemblies.

## Supporting information

Supplementary Protocol

