## Supplementary Protocol for "A Simple Enhancement for Gibson Isothermal Assembly"

**Supplementary Information**

**Protocol for the Purification of Recombinant *Tth* DNA Ligase**

The *ligA* gene from *Thermus thermophilus* (*Tth*) HB8 genomic DNA (Takara 3071) was amplified using nested PCR. It was cloned it into AsiSI (NEB R0630) and PmeI (NEB R0560) digested pFN18A HaloTag T7 Flexi Vector (Promega G275A) using our homemade Gibson Assembly Mix and transformed into Single Step (KRX) Competent Cells (Promega L3002) for protein production.

All solutions for protein purification were made with UltraPure water (ThermoFisher Scientific 10977015) and filtered through a 0.45 µm filter (Corning 430514).

Purified *Tth* DNA ligase was grown and purified using the HaloTag Protein Purification System (Promega G6270) according the manufacturer protocols with some deviations as follows. A single fresh colony was grown in a 5 ml LB liquid culture for 6 hours at 37C with 0.4% glucose (Millipore Sigma G7021). 1 ml of this culture was then added to 100 ml of LB with 0.05% glucose and 0.1% rhamnose (Promega L5701) and grown for 20 hours overnight at 25C before pelleting.

The pellet was resuspended in the recommended purification buffer with 0.005% IGEPAL Ca-630 (Millipore Sigma I8896) and 1 mM DTT (Goldbio DTT10). To lyse cells, 556 µl of 10x FastBreak Cell Lysis Reagent (Promega V8571) was added as well as the recommended RQ1 DNase (Promega M6101) and lysozyme (Millipore Sigma L2879). 50 µl of a SIGMAFAST Protease Inhibitor Tablet (Millipore Sigma S8820) dissolved in 100 ml of water was also added. This mixture was agitated on a shaker for 15 minutes at room temperature to allow for complete lysis. The lysate was then diluted to 10 ml with the original purification buffer above, then the recommended amounts of ATP (Millipore Sigma A7699) and MgCl_2_ (Millipore Sigma 442611) were added, and the lysate incubated at 37C for 10 minutes. The lysate was then cleared by centrifugation as recommended and to the cleared lysate EDTA (Millipore Sigma 324506) was added to 4 mM and DTT (Goldbio DTT10) was added to 20 mM for the binding step. The Batch/Column Combination Method from the HaloTag Protein Purification System (Promega G6270) protocol was used for the remainder of the purification. Washing with 60 ml of purification buffer with 0.5 mM EDTA was carried out to ensure complete removal of the lysis solution. Elution was done with single resin volumes, collecting individually and pooling based on the highest 280 nm absorbance (2 ml total).

The eluate was brought to 5 ml ensuring the final solution consisted of 50% glycerol, 10 mM Tris-HCl (pH 7.5, ThermoFisher Scientific 15567027), 50 mM KCl (ThermoFisher Scientific AM9640G), 1 mM DTT (Goldbio DTT10), and 0.2 mg/ml BSA (Millipore Sigma A9418). This purified enzyme was stored at -20C. The activity was determined with serial dilutions using the unit assay conditions supplied with commercially available *Taq* DNA ligase (NEB M0208) using the supplied buffer and λ DNA-BstEII Digest (NEB N3014) and one unit of commercially available *Taq* DNA ligase as a control (NEB M0208).
